# miR targetome of primary human keratinocytes reveals a function for non-conserved binding sites

**DOI:** 10.1101/2022.07.04.498673

**Authors:** Lalitha Thiagarajan, Jingqing Zhang, Sam Griffiths-Jones, Svitlana Kurinna

## Abstract

The homeostasis of the human body is protected by the skin, where the process of keratinocyte differentiation in outer layers has a crucial role. Cessation of proliferation in the basal layer of keratinocytes and initiation of their subrabasal functions are tightly controlled at the level of gene transcription and message translation. A subset of mRNAs has to be repressed during differentiation, and microRNAs are known to contribute to this by directly binding mRNAs at the 3’UTRs. Using results of RNA sequencing from human primary keratinocytes during induced differentiation, we evaluated the predicted binding of highly, moderately, and lowly expressed miRs to their target mRNAs. We found that moderately expressed miRs can regulate more mRNAs, and that they do so using both conserved and non-conserved canonical binding. The cumulative score for the majority of repressed mRNAs revealed a surprisingly weak binding to miRs, and we found a significant contribution of non-conserved sites to the repression of the targets. While the presence of at least one conserved site was necessary for the miR function, its weak binding may be reinforced by a non-conserved site. Together, we found that the combination of conserved and non-conserved sites lower the binding threshold for miR-mRNA interactions to assume a tighter repression of the mRNA target during cell differentiation.

## INTRODUCTION

Human skin protects homeostasis through the barrier function created by the outermost skin layer, the epidermis. Epidermis is a stratified epithelium differentiating from keratinocyte stem cells (KSC) contained in the basal layer, which divides to maintain the pool of keratinocytes attached the basal lamina as well as the supra basal cells (1). Differentiation of basal keratinocytes as they move upwards to create protective layers of epidermis is essential to maintain the integrity of the skin barrier. Keratinocyte differentiation is transcriptionally regulated by the epidermal differentiation complex (EDS) genes, acetylation of histone H3K27, and a complex interplay of well-defined transcription factors (TF), including p63, RANX2, and NRF2 (2-4). While TF-mediated, epigenetic mechanisms of basal-to-suprabasal transition and the skin barrier formation are extensively studied (5) much less is known about 1) mechanisms that maintain the basal, undifferentiated state of KSC and 2) the role of post-transcriptional mechanisms in differentiation of keratinocytes. As both of these questions are essential to understanding of skin repair and disease, in this study we aimed to elucidate the role of microRNAs (miRs) as the endogenous regulators of post-transcriptional regulation in basal keratinocytes during differentiation.

miRs are single stranded RNAs with a length of 19-25 nucleotides. There are 772 known human miR families, and a total of ∼40000 entries across animals and plants in the latest update of the miRBase database (6). miRs bind complementary sites of target mRNAs, usually in the 3’UTR, resulting in translational repression or degradation of the target mRNA (7). The primary determinant of targeting is 6-8nt at the 5’ end of the miRNA, called the ‘seed’ sequence. The majority of human genes may be regulated by miRs, and one miR can bind and regulate several target mRNAs, while one mRNA may also be regulated by a group of miRs (8). The complex regulatory network of miRNAs and their target mRNAs requires a combined experimental approach to investigate biological relevance (9).

While each miR is predicted to target hundreds of mRNAs, *in vivo* such interactions depend on the miR-mRNA pair actually ‘finding’ each other. This event depends on the levels of miR and mRNA expression and intercellular localisation, and may also depend on the cell type and the stage of cell and tissue development. Previously, we found that expression of the miR-29 family is limited by the degree of differentiation and the cell cycle stage of keratinocytes in human epidermis (4,10). In this study, we hypothesised that miRs specifically expressed in keratinocytes will bind the target mRNAs during early stages of differentiation and thus contribute to the regulation of gene expression observed during this process. We used small RNA and mRNA sequencing data from basal undifferentiated keratinocytes to predict miR-mRNA pairs and to follow the changes in target mRNA expression during *in vitro* induced differentiation of primary human keratinocytes (4). We found that while miRs may initially bind and repress target mRNAs, the majority of mRNAs change expression independently of miRs initially present in the cell. While this regulation can be specific to keratinocytes, it provides an example of selective function for miR *in vivo*, and emphasises the necessity of parallel sequencing and biochemical pull-down assays of small ncRNAs and mRNAs in global targetome studies.

## RESULTS

### Limited miRs function during differentiation of human keratinocytes

Publicly available small RNA-seq datasets were analysed to assess the expression of miRs in human primary keratinocytes isolated from foreskin and maintained in the ‘basal’ undifferentiated conditions according to the published differentiation protocol *in vitro* (1,2). From 8,858,015 reads across triplicate samples, 81.4% percent of reads mapped to 537 miRs that are found in miRBase (6) with average density ranging from 1 to 2,000,000 reads (Supplementary Table 1). Thus, approximately a quarter of known miRs recorded in miRbase (6) are expressed in basal human epidermis.

From 537 miRs, we selected 11 miRs representing groups of high, medium, and low levels of miR expression in keratinocytes, including the miR-122 with zero detected reads as a control (Table 1). A list of predicted mRNA targets for each miR was compared to the mRNAs expressed at the same level of keratinocyte differentiation (basal cells, D0). The differentiation model and the flow of the analysis is explained in Figure 1A and 1B, respectively. When primary cells isolated from human epidermis reach confluence, they start early differentiation at D1. This corresponds to the supra basal layer of cells in the skin and comprises a shift of epigenetic changes followed by the changes in transcription and translation at D1-D5 of incubation (Figure 1A). We hypothesised that miRs will bind their mRNA targets very early in this process, between D0 and D1, with most changes in mRNA repression manifesting at D3 and possibly, extending to D5. Thus, we predicted mRNA targets for miR expressed at D0 and repressed at D1-D5, and compared the lists of 100 top mRNA targets for 11 representative miRs (Figure 1B). We then analysed the cumulative weighted context score (CWCS), which determines the strength of miR-mRNA binding, for the top 100 predicted targets of the 11 chosen miRs (Figure 2A). By comparing the heat map of mRNA expression from D0 to D5 of differentiation, we expected a gradual distribution of repression for mRNA targets for each miR, manifested by the gradual loss of the colour on the heat map. This would indicate a spectrum of binding of miRs to their mRNA targets in undifferentiated keratinocytes, with highly expressed miRs perhaps binding and repressing more mRNAs even with low CWCS. While it was true for some miR-mRNA pairs, most mRNAs changed expression in a less gradual manner, with mRNAs displaying sharp rise and fall in the transcript levels seemingly independently of the paired miR (Figure 2A). Since a miR-mediated repression detected at D0 continue into at least early differentiation of keratinocytes (represented by D1-D3), this could result in both reversible and irreversible repression of mRNAs (3). miRs target mRNAs to degradation by the AGO proteins in the RNA-induced silencing complex RISC but may also withhold mRNAs from transcription by a transient sequestration in cytosol granules (11). Transcriptional regulation, on the other hand, can result in a more dynamic downregulation of mRNA levels via active repression or loss of activation at the promoter or enhancer (1,5). The expression changes of mRNA targets paired with 11 miR in human keratinocytes (Figure 2A) support both transcriptional and translational mechanisms. While the CWCS-based matching of miRs to mRNA targets may be used to predict their binding, most mRNAs are regulated in the miR-independent manner during differentiation of human keratinocytes.

**Table 1.** 11 selected miRs and their expressions.

| miR | chromosome | Start location | RPKM |
| --- | --- | --- | --- |
| hsa-miR-21 | chr17 | 57918634 | 1801378 |
| hsa-miR-205 | chr1 | 209605512 | 595121 |
| hsa-miR-182 | chr7 | 129410285 | 448593 |
| hsa-miR-92a-1 | chr13 | 92003616 | 170416 |
| hsa-miR-23a | chr19 | 13947383 | 102014 |
| hsa-miR-29a | chr7 | 130561506 | 68724 |
| hsa-miR-28 | chr3 | 188406622 | 36924 |
| hsa-miR-221 | chrX | 45605607 | 5106 |
| hsa-miR-34a | chr1 | 9211794 | 1259 |
| hsa-miR-302b | chr4 | 113569646 | 142 |
| hsa-miR-122 | chr18 | 56116306 | 0 |

**Figure 1.**
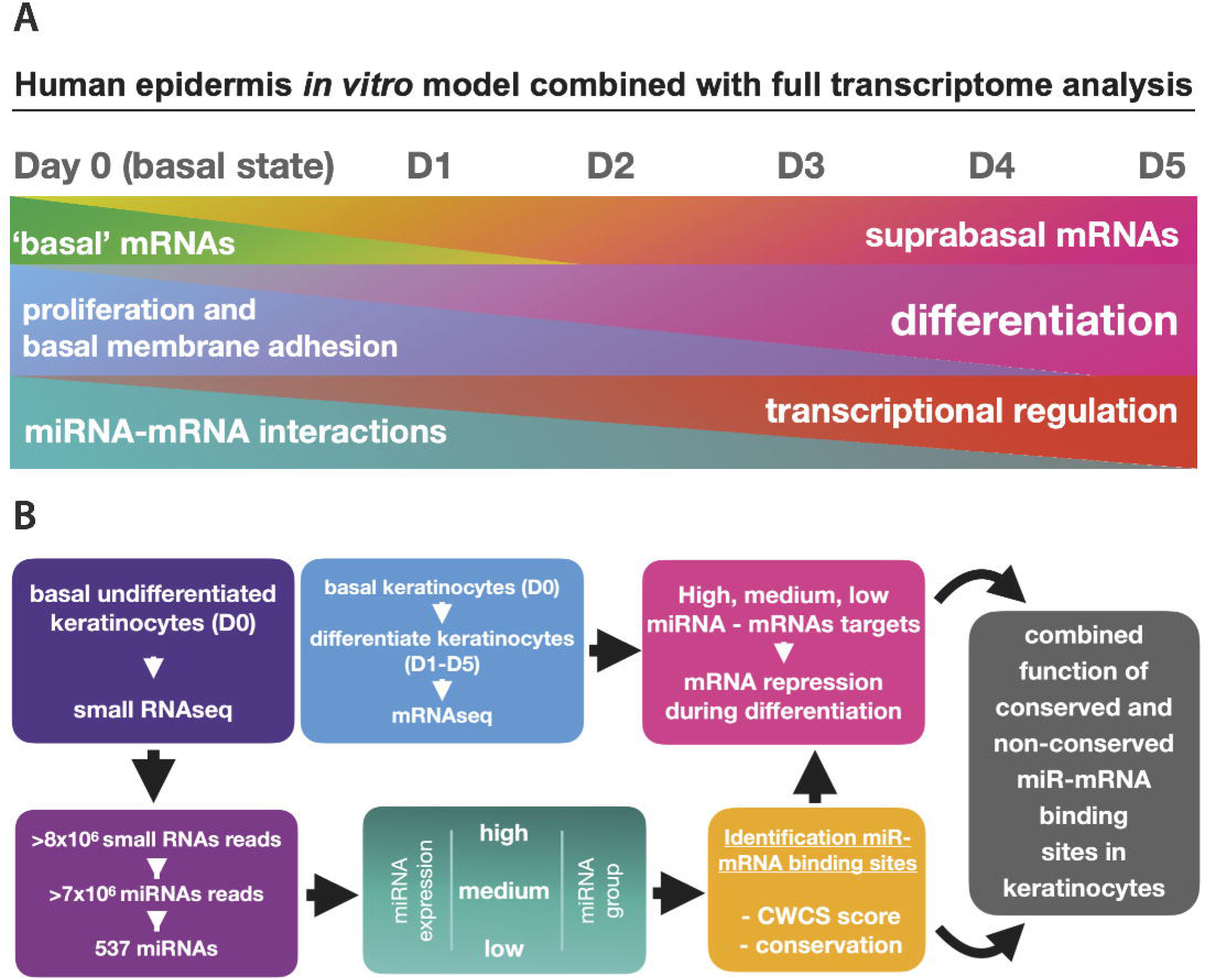
(A) Diagram of *in vitro* human keratinocyte differentiation model showing gradual changes in gene expression mediated by TF and miRs (B) Flowchart of miR and mRNA sequencing and basepair match analyses used in the paper.

**Figure 2.**
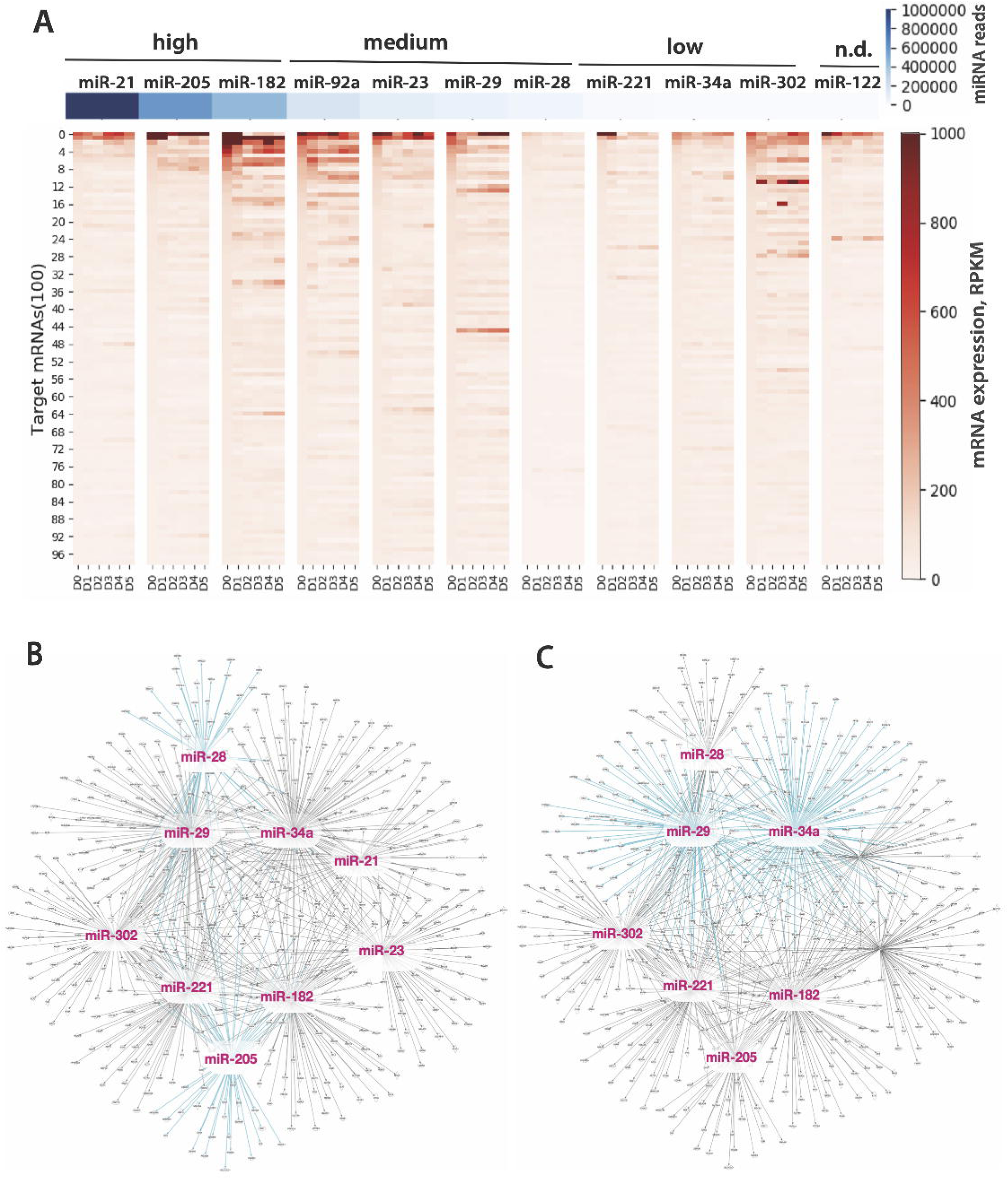
(A) Expression of selected miRs and their top 100 mRNA targets distributed along the cumulative weighted context score at day 0 of differentiation and changes in expression of 100 mRNAs targeted by 11 representative miRs at D0 to D5 of differentiation. (B) miR target network created using Ingenuity Pathway Analysis (IPA). Each ray-like line coming out of a candidate miR represents a target node predicted by IPA database and showing loss of expression in the RNAseq analysis.

Most surprisingly, the highly expressed miR-21, miR-205, and miR-182 did not steadily suppress the majority of the 100 predicted mRNA targets, with some of their targets showing a decline in RPKM after the onset of differentiation at D1 (Figure 2A). The expression change in mRNA targets of medium-expressed miR-92a, miR-23, and miR-29 showed very few mRNAs that decreased through the course of differentiation, suggesting an additional function for the medium-expressed miRs in keratinocytes. miR-28 may not have suppressed its targets as they show low levels of expression overall (Figure 2A). The abundant mRNAs in miR-221, miR-34a, and miR-302 from the low expressed group had changed irregularly, going from high to low RPKM during early differentiation and increasing again at D4 and D5. It is possible that most of targets of low expressed miRs are therefore regulated transcriptionally. The mRNAs of undetected miR-122 would be then entirely regulated through an miR-122-independent mechanisms (Figure 2A). Overall, these results suggest that only a limited number of predicted miR-mRNA pairs result in functional interactions *in vivo*.

### Non-abundant miRs have a wider targetome

To get insight into miR-dependant changes in expression of mRNAs, we used Ingenuity Pathway Analysis (IPA) to present the targetome of 11 miRs based on the IPA knowledge database (12). Two miRs that connected to a relatively low number of the interacting targets in the network were miR-28 and miR-205, both having medium to high expression in keratinocytes. While we expected a limited targetome for miR-28 based on a small changes in mRNA target expression and a weak CWCS, the IPA results for miR-205 function suggest that this miR binds to a few specific mRNAs in keratinocytes and does not regulate the rest of its predicted targetome (Figure 2B and 2C).

Interestingly, the miRs with the highest number of targets in the IPA-generated network were the medium and low expressed miR-29 and miR-34a (highlighted in Figure 2C). A closer look at the expression and binding score of individual miR-mRNA pairs suggests selectivity in targets. For example, one predicted target mRNA of miR-34a, RRAS, has the best CWCS among all predicted mRNAs, which means that miR-34a has the potential to strongly inhibit RRAS. The expression of only two mRNA, PDXK and UBE2G1, were lower than that of RRAS, which suggests that miR-34a plays a major role in suppressing of RRAS. PDXK and UBE2G1 do not have a good CWCS of interaction with miR-34a, which suggests that PDXK and UBE2G1 are regulated by a miR-34a independent mechanism (Supplementary Table 2). We hypothesised that the moderately expressed miRs may achieve a better regulatory potential through weaker, less conserved binding sites with high CWCS.

### Non-abundant miRs use non-conserved, weak binding sites

To test this, we used all predicted target mRNAs having a full range of CWCS of binding to the low- or medium-expressed miR ‘seed’. We found that non-abundant miRs showed significantly wider range of mRNA targets repression compared to mRNAs targets of abundant miRs (Figure 3A, quantified in Figure 3B). We chose medium expressed non-abundant miR-29 to compare the distribution of highly conserved miR-mRNA binding sites, as accessed by the CWCS. The analysis of miR-29 to target mRNAs binding in Targetscan (13) revealed that most mRNAs that are predicted to bind miR-29 through a 6-7-mer ‘seed’ match with low strength binding score ranging between 0 and -1, which we labeled as ‘very weak’ and ‘weak’ binding sites. Loss of mRNA reads was the only mechanism of miR-29-mediated regulation that we could test based on the RNAseq analysis, so we focussed on the of mRNAs that decreased expression during day 1, day 3, and day 5 of differentiation. We observed a strong shift of miR-29 targets along the CWCS axis with most target mRNAs grouping around very weak and weak binding (Figure 3C and Suppl. Table 3 for all miR-29 targets at day 1-5). We attributed this result to the presence of non-conserved sites, which contribute to the CWCS (13). It is known that the non-conserved sites, which outnumber the conserved sites 10 to 1, also contribute to miR-mediated repression (14). We hypothesised that the weak binding and repression of mRNAs in keratinocytes may be mediated by both conserved and non-conserved sites. Targets that were down-regulated either on day 3 or on day 5 of differentiation were pooled together for contingency analysis testing the contribution of conserved and non-conserved sites to the repression of mRNAs. While the presence of a non-conserved site was not required for the downregulation of the entire pool of targets with all scores, we suggested that the effect of non-conserved sites is masked by fewer repressed mRNAs with medium to strong CWCS (Figure 3D). Indeed, the repressed mRNAs with only one conserved site or with one conserved plus one poorly conserved site showed a significant contribution of non-conserved sites to the loss of target expression (Figure 3E). Thus, there’s a function for non-conserved sites but only in conjunction with the conserved sites, where they together contribute to the repression of the mRNAs.

**Figure 3.**
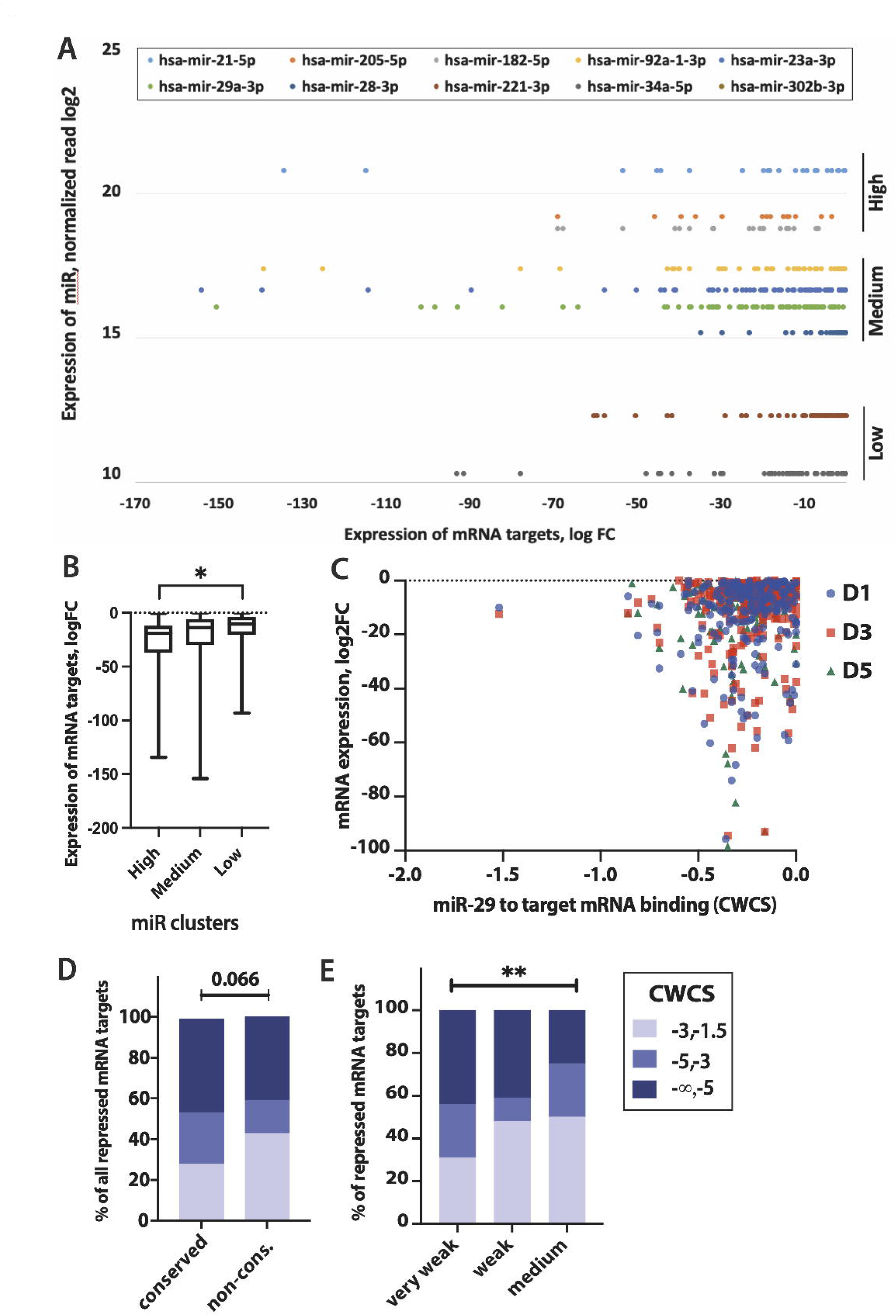
(A, B) Changes in expression in the predicted mRNA targets of 11 miRs during differentiation as a function of miR-mRNA binding cumulative weighted context score (CWCS). miRs are grouped as in Figure 2A. (C) Categories of CWCS shows grouping of miR-29 targets changing the expression in relation to the -5 logFC cut-off. (D, E) same cut-off applied and extended for all predicted miR-29 targets cumulatively changing the expression during differentiation. The contribution of conserved and non-conserved sites to the changes in target mRNA expression does not differ significantly in contingency analysis of the total context ++ score (TCS) from TargetScan but shows a significant contribution of both conserved and non-conserved sites to the changes in mRNA targets with TCS above -0.9.

To further test this, we plotted changes in mRNA expression from day 0 to day 3 and 5 of differentiation as a function of CWCS for 11 miRs. The binding score of nearly all mRNA targets of 11 miRs was in a range of weak, non-conserved sites (Supplementary Figure) (15). The predicted targets of miR-182, miR-92a, miR-23, and miR-29 showed more changes in expression compared to other miRs, suggesting that medium expression of miRs better positions them for the regulation of mRNAs in human keratinocytes.

## DISCUSSION

In this work, we used the data from the in vitro differential model of human keratinocytes analogous to the one utilised in previous studies of the role of TF and miR in differentiation (1,2,4). This is important because it provides the uniformity between the datasets obtained from ChIPseq and RNAseq analyses. Essentially, this study completes the profiling of primary human keratinocytes isolated from foreskin epidermis as a recognised consistent source of clonal KSC, transitionally amplifying ‘basal’ keratinocytes, and differentiated supra basal cells. Taking into account the previous work done using keratinocytes of the similar origin and consistency of in vitro differentiation, we can make the following conclusions. 1) While the transcriptional regulation is the strongest driver of differentiation both in the magnitude of the gene expression changes and the irreversibility of the following phenotypical change, miRs are important in mediating repression of mRNAs during keratinocyte differentiation; 2) miRs with the medium level of expression are most influential regulators of mRNA repression, and they do so via both ‘strong’ and ‘weak’ binding sites; 3) these miRs comprise the best candidates to target in skin repair and disease as the combination of ‘weak’ binding and relatively low expression of miRs allow the gradual and possibly, reversible therapeutic regulation of mRNA expression.

The transcriptional plasticity of KSC and immediate supra basal layers of human keratinocytes is attributed to both rapid and gradual changes of gene expression (16). Here, we focus on finding miR-mRNA pairs using a combination of small RNA and mRNA sequencing in primary human keratinocytes with prediction-based pairing of miR ‘seed’ sequence to the binding site in the 3’UTR of the mRNA. The basal proliferating layer of keratinocytes is responsible for skin regeneration, while suprabasal keratinocytes differentiate to create the layers of the epidermis, which makes the outer skin barrier and supports body homeostasis. We used this system to evaluate how the levels of expression of predicted miR-mRNA pairs reflect the changes in mRNA expression reflected during differentiation of primary human keratinocytes. Such knowledge of the actual and cell-specific targetome of miRs is crucial for understanding of normal homeostasis and disease (11).

We selected a very limited number of miRs for our analysis as we sought to test the prediction models for miR-mRNA interactions based on the RNAseq data from primary cells of the same type and differentiation stage. Another study found that expression of most miRs increases in human keratinocytes and confirmed miR-203 and miR-23b as markers of differentiation in human epidermis (17). Consistent with this study, we found miR-203 among high-to-medium expressed miRs (Supplementary Table 1). In undifferentiated keratinocytes we however detected another family member miR-23a, which can target the same mRNAs as miR-23b (18). It is interesting that the *in situ* hybridisation revealed near-exclusive expression of miR-23b transcripts in suprabasal cells of human epidermis (17), which may take over its family miR-23a function when keratinocytes move from basal to supra basal layers. Together, these results suggest that expression of miRs from the same family is tightly controlled during differentiation in human epidermis.

Our analysis of miR-mRNA interactions revealed a function for non-abundant miRs like miR-29 and miR-34a in human keratinocytes during normal differentiation. This is important because miR-34a is known to suppress proliferation of basal keratinocytes (19) and its expression is lost in skin and oral SCCs (20). Consistently with our analysis, miR-34a expression increases with keratinocyte differentiation (10). *In situ* hybridisation detects higher expression of miR-29 in human epidermis (4) also emphasising a function for this miR in regulation of normal cell growth and differentiation. We extensively analysed the distribution of binding scores of miR-29 in relation to the degree of suppression of target mRNAs and found that the majority of down regulated mRNAs have a weak binding to miR-29. Importantly, both conserved and non-conserved sites correlated with the repression of target mRNAs during the time course of keratinocyte differentiation. While the presence of a conserved site seems to be a minimum requirement for predicted binding and miR-mediated repression, the selective depletion of seed-matching sites in mRNAs that are highly expressed in the same tissues as the miRs implies frequent non-conserved targeting (14). Thus, mRNAs that have a combination of conserved and non-conserved miR sites can be repressed even when the score of mRNA-miR interactions predicts a very weak binding. Taken together, these results demonstrate that medium-to-low expressed miRs may target more mRNAs through a more frequent, transitional, and weaker binding.

In this study, we used Targetscan based miR-mRNA cumulative binding score, but the results of miRanda-based binding analysis using mirSVR score also identified a significant number of non-canonical and non-conserved sites (21). mirSVR machine-learning method confirmed that conservation should be used in combination with other informative features to score target sites and not as hard filter (21). We conclude that non-conserved binding sites of the miR targetome in primary human keratinocytes contribute to the regulation of normal differentiation in the epidermis, a process essential for the homeostasis.

## METHODS

### Selection and analysis of RNAseq datasets

miR expression was analysed using GEO dataset GSM1127113, which used undifferentiated (D0) primary human foreskin keratinocytes as a source of small RNA isolated and sequenced on Illumina HiSeq 2000. The mRNA expression from undifferentiated (D0) and differentiated keratinocytes were analysed using the RNAseq GSE73305, where the same protocol was applied to isolate and maintain primary foreskin keratinocytes at D0, followed by a five-day course of differentiation (D1, D2, D3, D4, and D5). The mRNA expression was expressed in reads per kb of transcript per million (RPKM). The raw count files were normalised and converted into bigwig format with Galaxy (https://usegalaxy.org/) and visualised using integrative genomics viewer IGV. miRs were mapped to human genome hg19 using chromosomal coordinates for *H. sapiens* miRs from miRBase (ftp://hsa-mirbase.org/pub/hsa-mirbase/19/genomes/hsa.gff3). The output table file with the information of miR name, chromosome, start position, and normalised read counts are stored in Supplementary Table 2. Expression heat maps were generated using the read density in the logarithmic scale at the base 2.

### Analysis of predicted target mRNA expression for selected miRs

11 miRs were selected from three different expression groups of all mapped miR. The predicted target mRNAs of each miR was identified in TargetScan (http://www.targetscan.org/vert_72/), and sorted using the cumulative weighted context score (CWCS). The lower the CWCS, the stronger the predicted binding of a miR and its inhibition on target mRNA (13). Expression levels of predicted target mRNA from the RNAseq data for each of the 11 miRs were extracted from the RNA-seq data using a Python script. A complementary list of CWCS for each miR-mRNA pair was used to estimate the total inhibitory effect of miRs. Data in these files was sorted by expression levels of undifferentiated cells (D0) and top 100 mRNAs was used to represent the results for day 1-5 of differentiation. To create a visualisation for mRNA network of individual miRs, the data was analysed with <u>Ingenuity Pathway Analysis</u> (IPA) (12).

### Binding score and mRNA expression contingency analysis

Binding sites for a larger dataset of mRNAs were analysed using Targetscan (13), which divided them between highly conserved 7- and 8-mer sites with CWCS below or close to -1 and non-conserved sites mostly with 7 nt match between miR and mRNA 3’UTR.

Contingency plots of the down-regulated targets with only one conserved site or with one conserved plus one non-conserved site were generated using Prism by categorising total context++ score based on their strength of binding (13) and by comparing with the percentage of down-regulated mRNA targets in each of those categories. Correlational significance was tested by performing Chi-square test.

## Acknowledgements

The authors thank Dr. Poojitha Rajasekar, the University of Nottingham, for her help with the analysis of the distribution of predicted miR-29 targets.

## Authors’ contributions

L.T. assisted in developing of the project, performed miR-mRNA target score and functional analyses, and assisted in wring the manuscript. J.Z. developed and utilised Python codes for mRNA and miR sequencing analyses; S. G.-J. participated in designing and discussing the project and revision of the manuscript. S.K. designed the project, wrote the manuscript, and acquired funding. All authors made important comments on the manuscript.

---

**Supplementary Figure.**
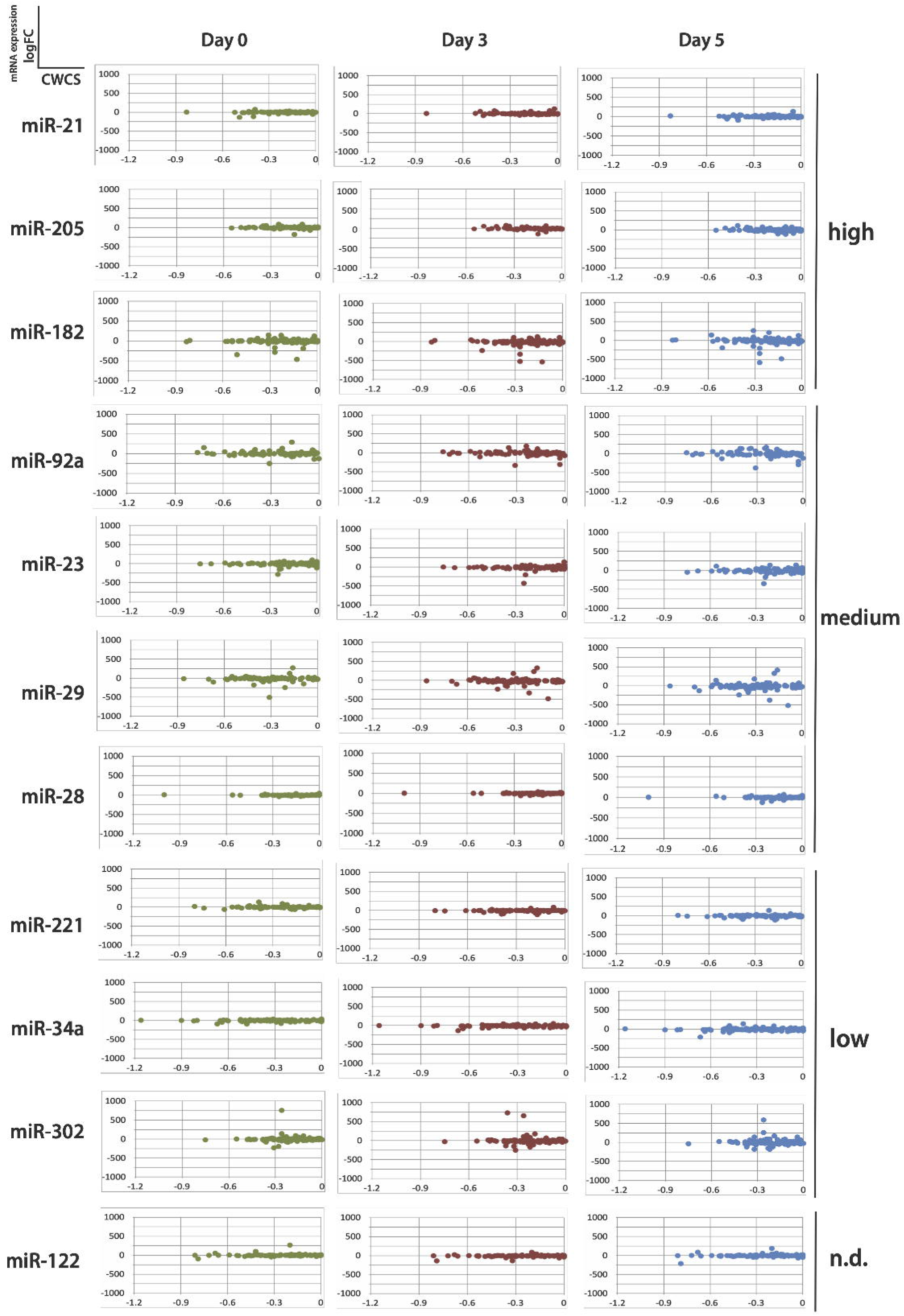
Changes in expression in the predicted mRNA targets of 11 miRs during day 0-5 of differentiation as a function of miR-mRNA binding cumulative weighted context score (CWCS).

